# Modeling the Structural Origins of Drug Resistance to Isoniazid via key Mutations in *Mycobacterium tuberculosis* Catalase-Peroxidase, KatG

**DOI:** 10.1101/230482

**Authors:** Matthew W. Marney, Robert P. Metzger, David Hecht, Faramarz Valafar

**Affiliations:** Department of Chemistry and Biochemistry, San Diego State University, San Diego, CA 92182-1030; Department of Chemistry, Southwestern College, Chula Vista, CA 91910; Biomedical Informatics Research Center, San Diego State University, San Diego, CA 92182-7720

## Abstract

WHO reported 10.4 million new tuberculosis (TB) cases and 1.8 million deaths in 2015, making *M. tuberculosis* the most successful human pathogen with highest mortality among infectious diseases.[1,2] Drug-resistant TB is a major threat to global TB control.[2,3] Recently Torres et al. [4] identified 14 novel substitutions in *M. tuberculosis-KatG* (the enzyme associated with resistance to isoniazid—an important first-line anti-TB drug) and demonstrated that 12 of the 14 can cause INH-resistance in *M. smegmatis*. This study presents an *in silico* structure-based analysis of these 14 amino acid substitutions using homology models and x-ray crystal structures (when available) in *M. tuberculosis*. Our models demonstrate that several of these mutations cluster around three openings in the KatG tertiary structure which appear to initiate channels to the heme group at the catalytic center of the enzyme. We studied the effects of these mutations on the tertiary structure of KatG, focusing on conformational changes in the three channels in the protein structure. Our results suggest that the 14 novel mutations sufficiently restrict one or more of these access channels, thus potentially preventing INH from reaching the catalytic heme. These observations provide valuable insights into the structure-based origins of INH resistance and provide testable hypotheses for future experimental studies.

## 1 Introduction

Drug resistant strains of *Mycobacterium tuberculosis* (*M. tuberculosis*) pose a continuously evolving threat to human health worldwide. It is estimated that nearly one fourth of the world’s population is infected by latent *M. tuberculosis*.[5] In 2015, the World Health Organization (WHO) reported 10.4 million new active cases and 1.8 million deaths from tuberculosis (TB).[2] Multi drug-resistant (MDR) strains, resistant to the two most powerful first line anti-tubercular drugs isoniazid (INH) and rifampicin (RIF), comprised greater than 20% of existing and greater than 7% of new cases in 2015.[2,3] Attempts to develop replacements for INH and RIF have not been encouraging, generally yielding drug candidates less potent and/or more toxic than INH and RIF.[6] Efforts to better understand the mechanisms by which mutations render INH and RIF ineffective are prerequisite to development of successful new treatments for TB and accurate diagnostics.[7,8] In this paper, we focus on those mutations that cause INH resistance in *M. tuberculosis.*

INH is a pro-drug. Once taken up by *M. tuberculosis*, INH serves as a substrate, along with NAD^+^, for the catalase-peroxidase enzyme KatG-catalyzed formation of nitric oxide (NO) and isonicotinyl-NAD.[9–11] (Figure 1) Isonicotinyl-NAD binds to the active site of enoyl acyl carrier protein reductase, blocking fatty acid synthesis in general and the synthesis of mycolic acids, which are components of the *M. tuberculosis* cell wall, in particular.[10]

**Figure 1.**
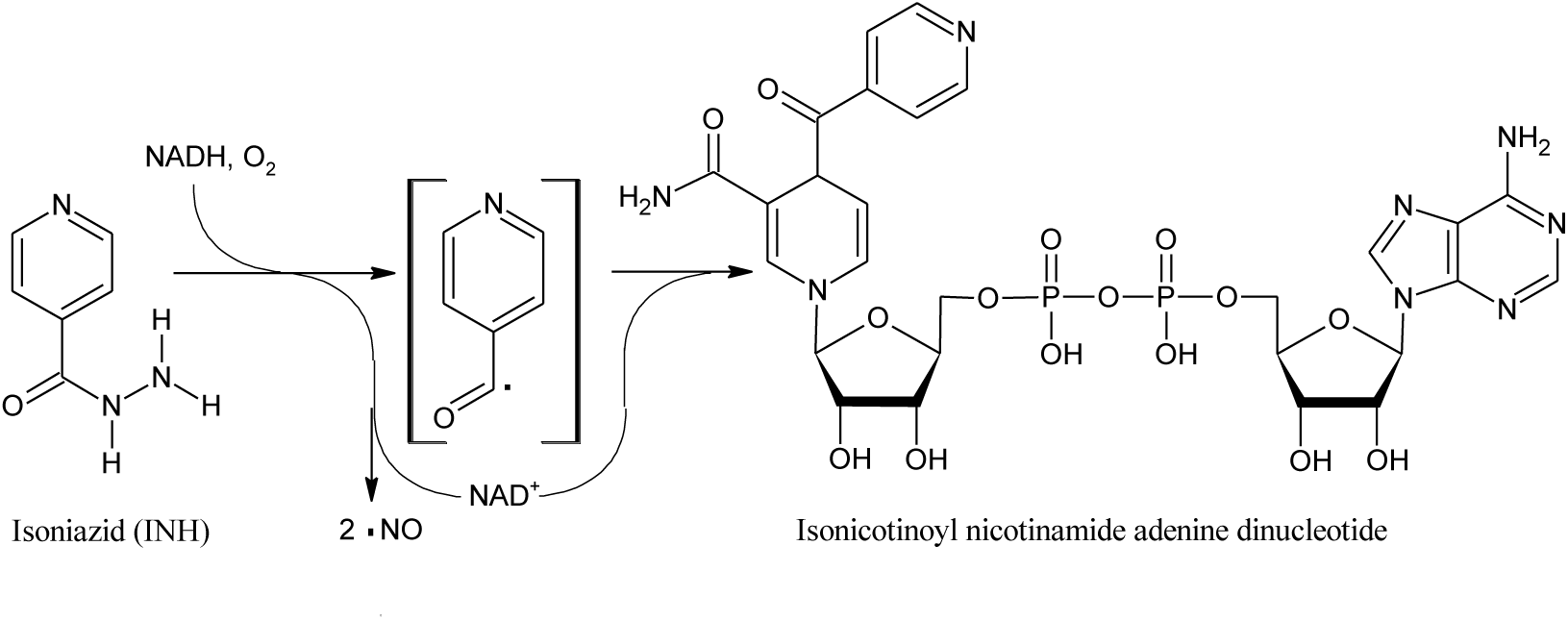
The structure of INH, also known as isonicotynylhydrazide or pyridine-4-carbohydrazide, along with a schematic version of the reaction of INH catalyzed by the catalase-peroxidase enzyme KatG based on findings of Singh *et al* [9] and Timmons *et al*. [10] The free radical nature of the reaction and formation of NO is experimentally well-established[9,11] but not all intermediates may have been identified and the radical shown in brackets, while possibly a reaction intermediate, is for illustrative purposes only. Thus, no attempt is made to show a balanced reaction. The product isonicotinoyl nicotinamide adenine dinucleotide binds at the active site of the enzyme enoyl-acyl carrier protein reductase, stopping *M. tuberculosis* fatty acid synthesis, including the synthesis of the cell wall component myconic acids.[10].

Resistance to INH is primarily caused by key mutations of the catalase-peroxidase, KatG, and/or promoter mutations in the *inhA* gene.[12,13] The most frequently observed mutation involving an amino acid substitution conferring INH resistance (KatG S315T) is believed to restrict a pathway into a catalytic heme center in the active site.[14—21]

In a recent molecular dynamics study, Pimentel and coworkers demonstrated that single amino acid substitutions in *Mtb*-KatG identified in INH-resistant clinical isolates from Brazil, can decrease the volume of the catalytic cavity as well as potentially alter the positioning of the catalytic heme.[22] The majority of amino acid substitutions studied were either in direct contact with the catalytic heme (S315T, S315R, S315I, S315N, S315G and G273C) or were located in close proximity, within 3 Å (P232R). They were also able to demonstrate that amino acid substitutions located > 10 Å from the catalytic heme (A109V as well as the double mutant H97R, L200Q) can decrease the volume of the catalytic center.

Recently Torres et al. identified 14 novel amino acid substitutions in *M. tuberculosis*-KatG and, through mutagenesis, demonstrated that 12 of them cause INH resistance in cultures of *Mycobacterium smegmatis*.[4] Interestingly, these 14 different amino acid substitutions are distributed throughout the protein sequence making it very difficult to understand how these mutations, most being far from the 315 position, can lead to INH resistance. With this goal in mind, we have performed molecular modeling in order to understand the structure-based origins of INH resistance for these variant KatG proteins.

Kamachi et al. recently published an x-ray structure of WT type *Synechococcus elongates* KatG (*Se*-KatG) co-crystalized with three bound INH molecules (3WXO.pdb).[15] *Se*-KatG shares 55% sequence identity with the x-ray structure of wild type (WT) *M. tuberculosis*-KatG (2CCA.pdb).[23] The three binding sites of INH in the x-ray crystal structure 3WXO.pdb are believed to be along three different passages through which various substrates, including INH, could pass from the protein surface to the catalytic heme group.[23] The tertiary structure and overall fold of both protein domains are very similar, and a structural-based superposition of the two structures, shown in Figure 2, results in a virtual overlap of the three channels as well as the catalytic center with negligible RMSD value of 1.03 Å (including all atoms between the two x-ray structures).

**Figure 2.**
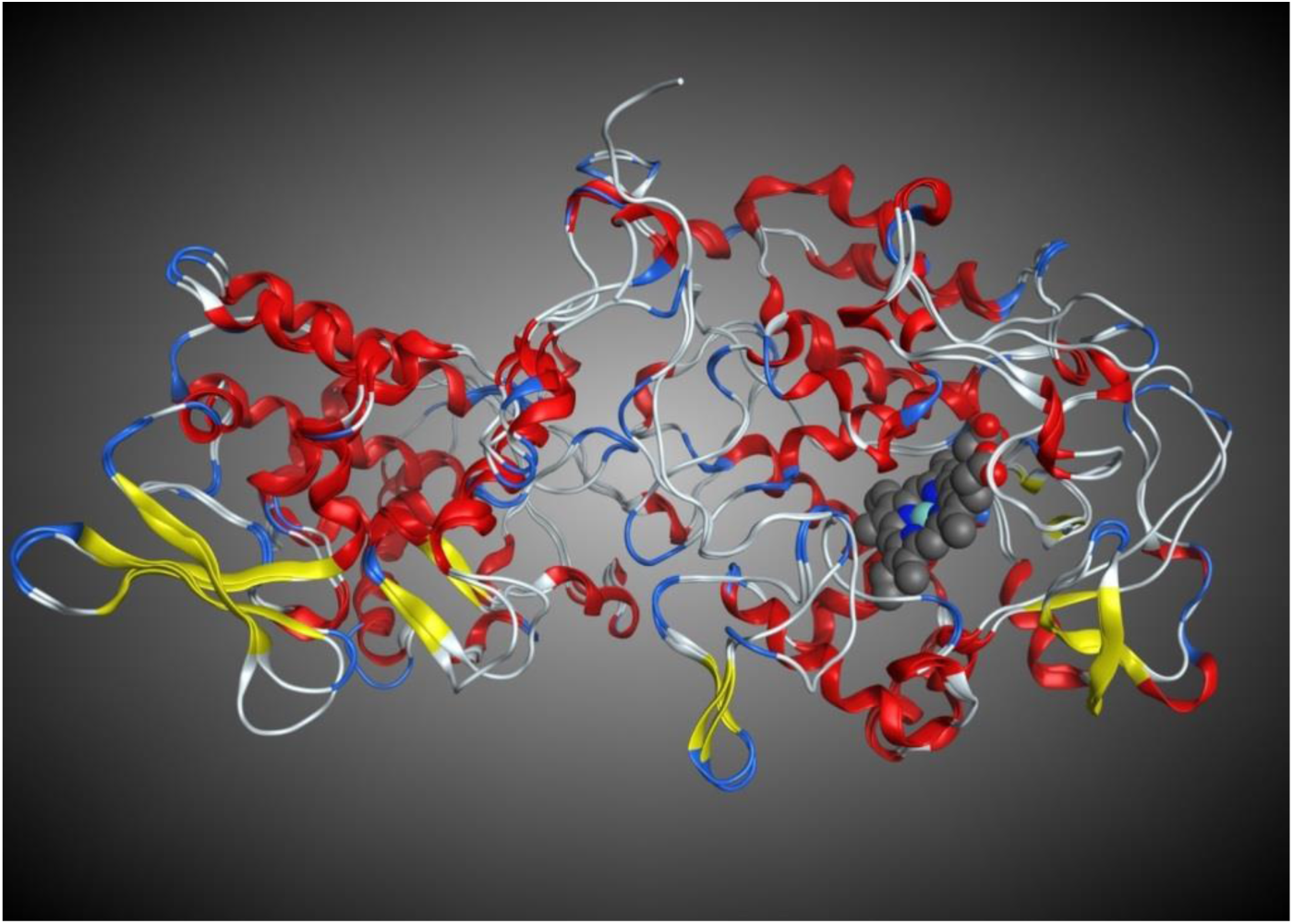
Structural superposition of the x-ray crystal structure of WT *M. tuberculosis*-KatG (2CCA.pdb) with WT *Se*-KatG (3WXO.pdb) illustrating the similar secondary and tertiary structures, overall 1.03 Å *RMSD*. The heme from 2CCA.pdb is shown in blue and grey.

Using *M. tuberculosis*-KatG 2CCA.pdb as a template, we generated homology models of the 14 variant KatG sequences reported by Torres et al.[4] Several of these amino acid substitutions appear to cluster around the three openings in the KatG tertiary structure that lead directly to the heme group at the enzyme’s catalytic center, the same 3 sites that were found in the x-ray structure of *Se*-KatG. [15] In this article, we report the results of our investigation into the structural impact of these 14 novel mutations on the tertiary structure of *M. tuberculosis*-KatG, focusing on conformational changes occurring in all three potential entry channels to the active site with mutant enzymes showing reductions in channel cross-section ranging up to ~40%. These observations provide valuable insights into the structure-based origins of INH resistance and provide testable hypotheses for future experimental studies.

## 2 Materials and Methods

### *2.1* Sequence Variation and Homology Modeling

The x-ray crystal structure of WT *M. tuberculosis*-KatG (2CCA.pdb) was downloaded from the RCSB PDB and imported into the comprehensive bio/cheminformatics software package MOE (www.chemcomp.com).[24] Using the sequence editor in MOE, variant sequences of KatG were generated, each with one of the 14 INH resistance conveying amino acid substitutions identified by Torres et al.[4] Homology models were then generated, with default MOE settings, for each mutant sequence using the x-ray crystal structure 2CCA.pdb as a template. *RMSD* values for each homology model from the template x-ray crystal structure, 2CCA.pdb, fell within the range of 0.25 to 0.39 Å.

### *2.2* Controls

Three thousand nine hundred and forty two sequences of *M. tuberculosis*-KatG were downloaded from the NCBI Protein Database (http://www.ncbi.nlm.nih.gov/protein). The set was reduced to ~100 non-redundant sequences that were observed in INH susceptible isolates. Reduction criteria included the exclusion of all duplicates (100% sequence identity), all WT, and all sequences harboring anything other than a single substitution. From this set, four sequences were selected as control. The intention was to demonstrate that our models could predict that the mutations do not confer resistance by predicting minimal steric effects. The four substitutions were: A110V [25], G316S [26], L499M [25], and L587P [26]. The lineage marker R463L which is broadly observed in susceptible and resistance isolates, and does not confer resistance, was also added to this list as the fifth control substitution.[4]

Homology models were then generated for each sequence, with default MOE settings and using the x-ray crystal structure 2CCA.pdb as a template. *RMSD* values for each homology model from the template x-ray crystal structure, 2CCA.pdb, fell within the range of 0.25 to 0. 39Å reported above. Each homology model was added to the structure-based alignment and measurements were performed on each of the channels. (Supplementary Tables ST1 and ST2)

### *2.3* Structure-Based Alignment

The x-ray crystal structure of WT *Synchococcus elongates*, *Se*-KatG (3WXO.pdb), containing the structure of bound INH was downloaded from the RCSB PDB and imported into MOE. Using the sequence and structural alignment functions *Align* and *Superpose* in MOE, all 14 variant *M. tuberculosis*-KatG homology models as well as 3WXO.pdb were structurally superimposed on 2CCA.pdb.

### *2.4* Mapping the Access Channels on M. tuberculosis-KatG

WT *Se*-KatG (3WXO.pdb) and WT-*M. tuberculosis*-KatG (2CCA.pdb) share 55% sequence identity as well as a highly conserved fold with almost identical secondary and tertiary structures, as indicated by the value of 1.03Å *RMSD*. (Figure 2) Using the structural superposition, the amino acids comprising the three access channels for potential INH entry to the catalytic center labelled 1, 2 and 3, were then mapped onto corresponding residues in the *M. tuberculosis*-KatG structure-based alignment including the 14 KatG amino acid variants identified by Torres et al.[4] These were highlighted yellow, red, and blue, respectively. (Figure 3)

**Figure 3.**
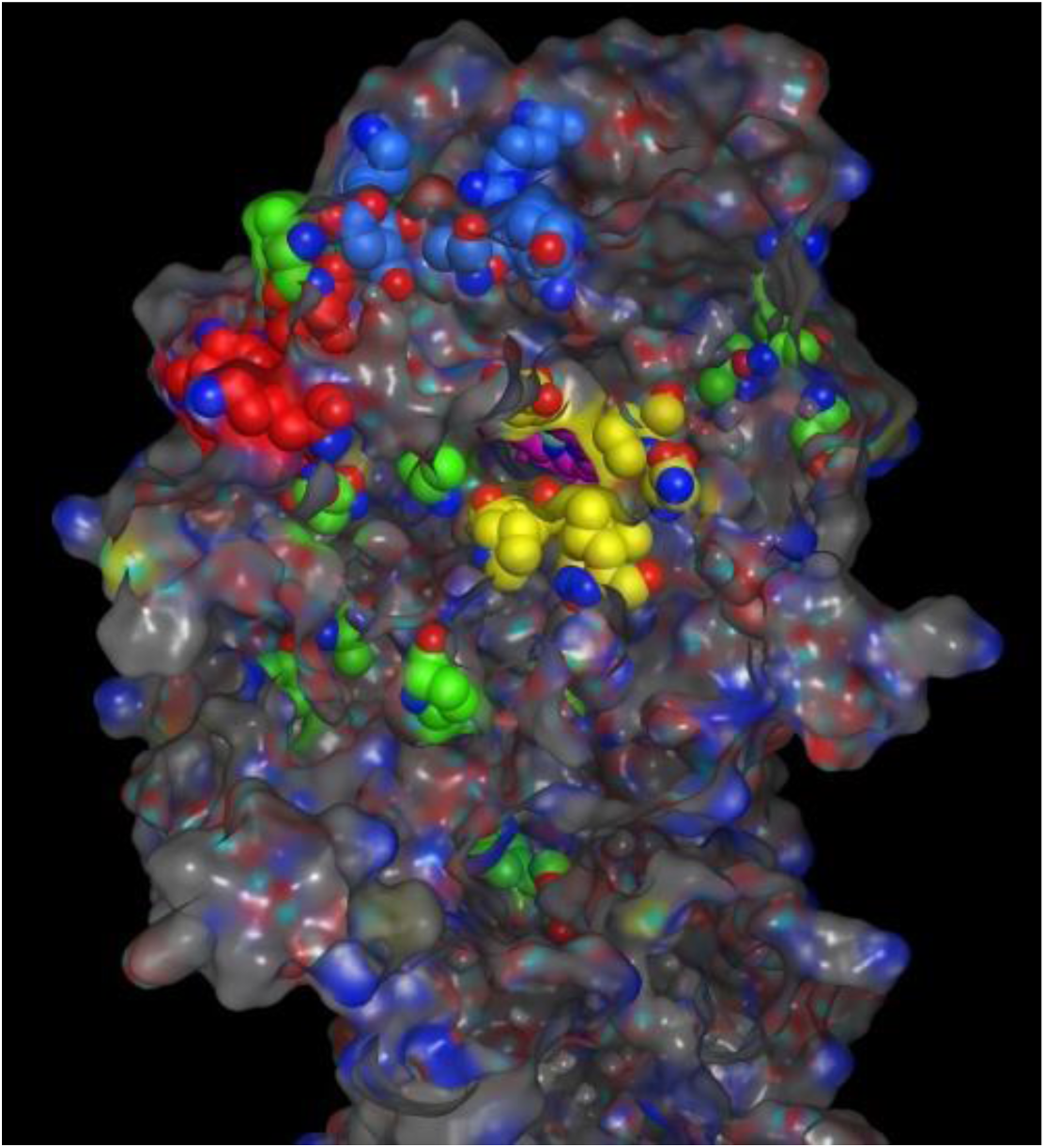
(Right) The x-ray crystalstructure of WT *M. tuberculosis*-KatG (2CCA.pdb) with Channel 1, 2 and 3 residues colored yellow, red, and blue respectively. The 14 resistance conveying amino acid replacements are colored green. The catalytic center containing the pink colored heme atoms, is visible through channel 1.

**Figure 4.**
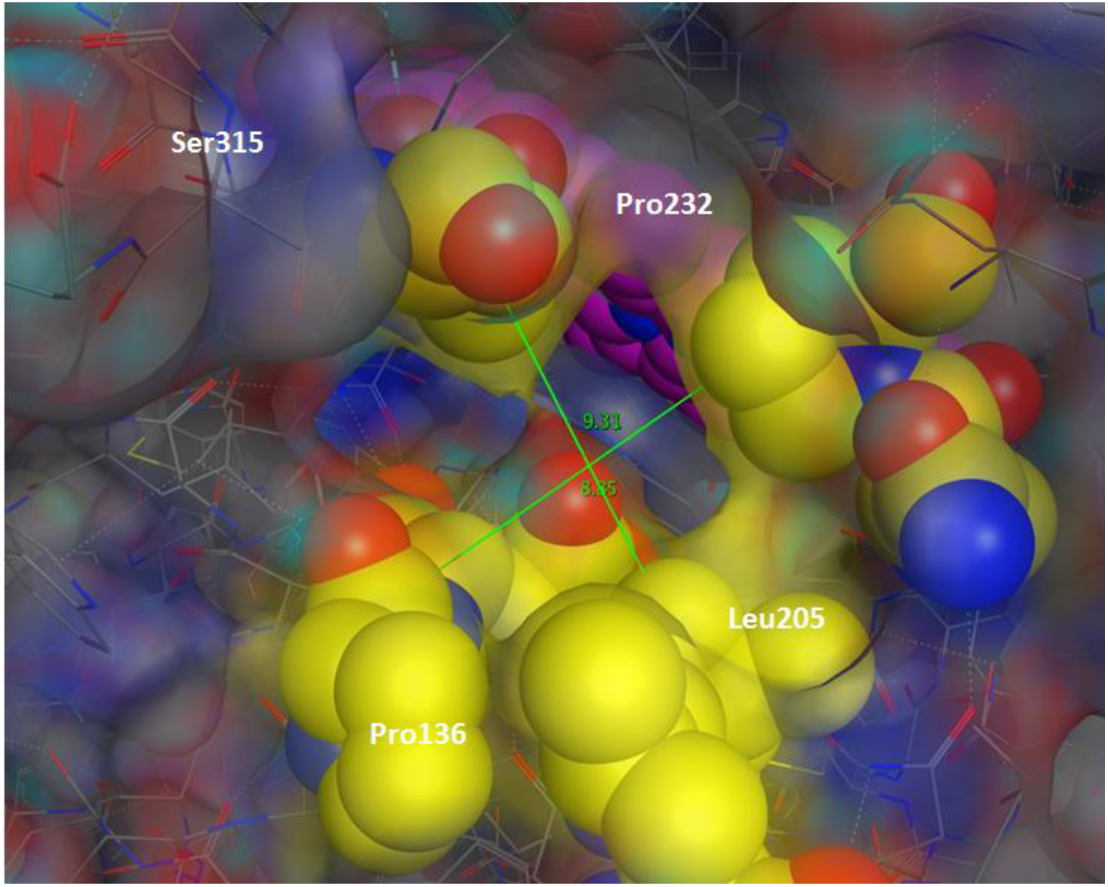
(Above) Channel 1 residues (in yellow) from WT *M. tuberculosis*-KatG (2CCA.pdb) superimposed on the x-ray structure of WT *Se*-KatG with bound INH (3WXO.pdb) in CPK coloring.

**Figure 5.**
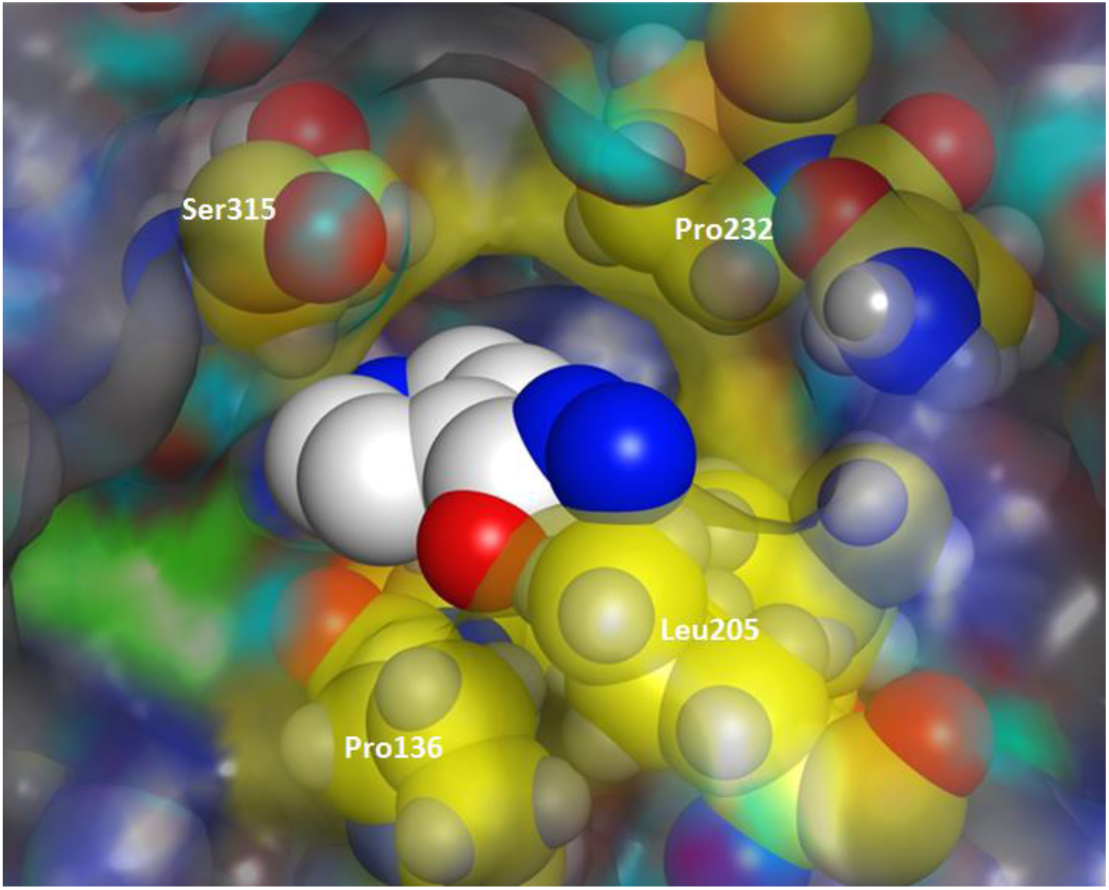
(Right) Illustration of the major and minor axis measurements on channel 1. The catalytic center (colored pink) visible through the channel (colored yellow). Major and Minor axis measurements made on all three channels in each of the KatG structures. These measurements were used to calculate the cross-sectional area presented in Table 1.

**Table 1.** Percentage Change in cross-sectional areas at the surface entrance of each of the three channels leading to the catalytic center with respect to corresponding sites in the WT x-ray structure, 2CCA.pdb as a result of the listed mutations. “>10 Å” indicates that the mutation was more than 10 Å away from all three channels. MIC values are those reported by Torres *et al.* [4] MIC^+^ values are estimations used in our analysis. (Please see the Method Section)

| KatG Mutation | Closest Channel | Site 1 % Δ Area (Å<sup>2</sup>) | Site 2 % Δ Area (Å<sup>2</sup>) | Site 3 % Δ Area (Å<sup>2</sup>) | MIC | MIC<sup>+</sup> |
| --- | --- | --- | --- | --- | --- | --- |
| WT | NA | 0% | 0% | 0% |  | 0.01 |
| S315T | Site 1 | -2.87% | -5.32% | -2.11% |  | 2.5 |
| **Novel Mutants reported by Torres et al.[4]** |  |  |  |  |  |  |
| Y64S | >10 Å | -19.74% | -20.82% | -19.54% | 3 | 3 |
| Y95C | Site 3 | -27.99% | -20.77% | -44.57% | 0.5 | 0.5 |
| P131T | Site 1 | -17.52% | -24.32% | -18.26% | 0.5 | 0.5 |
| A139P | Site 1 | -15.47% | -34.49% | -46.25% | >10 | 15 |
| D142G | Site 1 | -26.26% | -18.52% | -42.56% | >10 | 15 |
| A162V | >10 Å | -22.87% | -36.47% | -19.55% | 3 | 3 |
| G269D | >10 Å | -16.51% | -37.17% | -22.05% | >10 | 15 |
| T306P | Site 3 | -19.17% | -17.98% | -40.95% | ND |  |
| R385W* | >10 Å | -12.08% | -26.58% | -10.96% | >10 | 15 |
| D387G* | >10 Å | -22.46% | -23.35% | -32.06% | ≤10 | 8 |
| T394M | >10 Å | -16.47% | -32.83% | -33.69% | ND |  |
| Q439P | >10 Å | -8.48% | -36.28% | -27.94% | ND |  |
| F483L | >10 Å | -24.15% | -25.83% | -37.58% | 3 | 3 |
| A541D | >10 Å | -23.45% | -36.01% | -35.17% | >10 | 15 |
| **Nonresistance Conferring Controls** |  |  |  |  |  |  |
| A110V | >10 Å | +2.29% | +0.31% | -0.98% |  | 0.01 |
| L499M | >10 Å | +1.42% | +0.09% | +2.12 |  | 0.01 |
| L587P | >10 Å | -3.13% | +4.80% | -2.26% |  | 0.01 |
| R463L | >10 Å | -0.87% | +0.85% | +4.43% |  | 0.01 |
| G316S | Site 1 | -5.26% | +3.87% | +2.22% |  | 0.01 |

\* *M. smegmatis* mutants with these mutations did not grow in presence of INH (Torres et al.[4])

### *2.5* Cross-Sectional Area Measurements

Cross-sectional diameters of the surface opening of each channel were measured in the homology models of the canonical mutation S315T, the 14 novel variants, five variants observed in susceptible isolates, and of the WT KatG. (Supplementary Table ST1, Table 1, Figures 4 and 5). A second set of cross-sectional measurements were performed near the heme end of each channel. (Supplementary Table ST2, Table 2) The approximate cross-sectional areas for each of the homology models and the corresponding positions in the WT structure were calculated using equation (1) and compared to the WT.

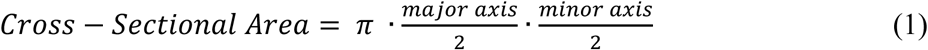

**Table 2.** Percentage Change in cross-sectional areas near the entrance to the catalytic center for each of the three channels with respect to corresponding sites in the WT x-ray structure, 2CCA.pdb. “>10 Å” indicates that the mutation was more than 10 Å away from all three channels. MIC values are those reported by Torres *et al.* [4] MIC^+^ values are estimations used in our analysis. (See Method Section)

| KatG Mutation | Closest Channel | Site 1 % Δ Area (Å<sup>2</sup>) | Site 2 % Δ Area (Å<sup>2</sup>) | Site 3 % Δ Area (Å<sup>2</sup>) | MIC | MIC<sup>+</sup> |
| --- | --- | --- | --- | --- | --- | --- |
| WT | NA | 0% | 0% | 0% |  | 0.01 |
| S315T | Site 1 | -22.11% | +8.99% | +9.51% |  | 2.5 |
| **Novel Mutants reported by Torres *et al* [4]** |  |  |  |  |  |  |
| Y64S | >10 Å | -1.44% | -14.00% | +66.27% | 3 | 3 |
| Y95C | Site 3 | -3.12% | -20.62% | +27.72% | 0.5 | 0.5 |
| P131T | Site 1 | -1.46% | -14.67% | +65.43% | 0.5 | 0.5 |
| A139P | Site 1 | +25.77% | -21.06% | +26.96% | >10 | 15 |
| D142G | Site 1 | +2.29% | -18.17% | +46.88% | >10 | 15 |
| A162V | >10 Å | -6.95% | -10.39% | +54.81% | 3 | 3 |
| G269D | >10 Å | +3.75% | -13.92% | +49.39% | >10 | 15 |
| T306P | Site 3 | -0.80% | -15.84% | +59.09% | ND |  |
| R385W* | >10 Å | +1.81% | -17.59% | +52.97% | >10 | 15 |
| D387G* | >10 Å | -2.57% | -19.94% | +51.70% | ≤10 | 8 |
| T394M | >10 Å | -0.92% | -10.42% | +63.22% | ND |  |
| Q439P | >10 Å | +15.90% | -9.85% | +72.80% | ND |  |
| F483L | >10 Å | -3.84% | -20.80% | +40.74% | 3 | 3 |
| A541D | >10 Å | -0.01% | -19.25% | +57.06% | >10 | 15 |
| **Nonresistance Conferring Controls** |  |  |  |  |  |  |
| A110V | >10 Å | +14.50% | -3.12% | +20.17% |  | 0.01 |
| L499M | >10 Å | +14.69% | -1.77% | +19.67 |  | 0.01 |
| L587P | >10 Å | +9.98% | +0.08% | +20.69% |  | 0.01 |
| R463L | >10 Å | +13.25% | -2.10% | +31.32% |  | 0.01 |
| G316S | Site 1 | +12.11% | -4.65% | +34.40% |  | 0.01 |

\* *M. smegmatis* mutants with these mutations did not grow in presence of INH (Torres et al.[4])

### *2.6* Correlation between Resistance Levels and Constriction of the Access Channels

In order to assess the potential role of steric effects on level of resistance, we estimated the correlation between Minimum Inhibitory Concentration (MIC) levels (reported by Torres et al. [4]) and the relative constriction of the channels with regards to WT-KatG. As mentioned before, two measurements were made in each channel of each model, one at the protein surface and one near the heme.

We studied the predictive power of five sets of measurements in predicting the reported MIC values. These measurements were: cross-sectional opening of the channels at the surface of the protein(*A_s_*), percentage change of*A_s_* as compared to the WT (%Δ*A_s_*), cross-sectional opening of the channels near the heme *A_h_*, percentage change of *A_h_* as compared to the WT (%Δ*A_h_*), and smallest cross-sectional area in each channel (between the surface and heme-end opening) *A_min_* = min(*A_s_*,*A_h_*).

In order to study the predictive power of the listed measurements and their relationship with MIC levels, we employed a variety of statistical, polynomial, rational, and exponential functions. Ninety eight different instances of the 18 different form functions listed in Supplementary Table ST3 were employed. Each of the five listed measurements from each channel was separately provided as input to the 98 functions. In all 1,470 different input-function combinations were tried. The outputs of the functions were used as estimates of the MIC values. We then studied the correlation between our estimates and the actual MIC measurements in order to identify the input-function combination that produced the most accurate MIC estimates.

We used both Pearson (*ρ_p_*) and Spearman’s Correlation Coefficient (*ρ_s_*) to estimate the level of correlation between the function outputs and the MIC values. [27] In all cases the relationship *ρ_s_* ≥ *ρ_p_* persisted. Since this is an indication of a monotonic nonlinear relationship, Spearman’s correlation is the more appropriate measure, and hence, in this manuscript, we only report *ρ_s_* values.

From the MIC ranges reported by Torres et al. [4] exact values were estimated using the following criteria: 1) for all isolates reported as having an MIC of greater than 10 *μg/ml*, 15 *μg/ml* was chosen as MIC. 2) 8 *μg/ml* was used for those isolates reported with an MIC level greater than 3 and less than 10 *μg/ml*. 3) For all susceptible control isolates with a KatG mutation or a WT KatG, we assumed the MIC value of 0.01 *μg/ml*—well below the WHO recommended MGIT960 cutoff (0.1 *μg/ml* [28]) for INH resistance. 4) For the isolate with the canonical KatG-S315T, we used the MIC value of 2.5 *μg/ml*. A wide range of MIC values have been reported in the literature for isolates with this canonical mutation. An MIC value of 2.5 *μg/ml* represents the average of these values. We also experimented by changing the estimated MIC values within the ranges reported by Torres et al. The effect on the correlation values was negligible for all input-function combinations. Tables 1 and 2 show the complete list of MIC values reported by Torres et al. [4] and those used in our experiments.

## 3 Results and Discussions

### *3.1* Homology Model Evaluation

The major assumption in this approach is that a single amino acid substitution will have minimal effect on the overall protein structure and function. In order to test the validity of this assumption, we downloaded all 48 x-ray crystal structures of KatG from six different species: *Burkholderia pseudomallei*, *Synechococcus elongatus*, *Mycobacterium tuberculosis*, *Haloarcula marismortui*, *Magnaporthe oryzae*, *and Escherichia coli*. The structures were imported into MOE and a structure-based alignment was performed (as described above in the Methods section). As illustrated in Supplementary Figure SF1, although these proteins have very different sequences, sharing only 13 sequence positions containing conserved amino acid residues, they all have essentially the same domain fold with overlapping secondary and tertiary structures, *RMSD =* 0.91Å. (Supplementary Figure SF1)

This analysis demonstrates that very different KatG sequences, coming from six evolutionary diverse archea, bacterial and eukaryotic species, all share essentially equivalent experimentally determined 3D structures. This result provides confidence in the use of homology models to model single amino acid substitutions. Each homology model generated was compared to the template x-ray crystal structure, 2CCA.pdb, with *RMSD* values ranging between 0.25 and 0.39Å, with the average *RMSD* being 0.31 ± 0.02 Å. Ramachandran plots were generated to identify models having unusual or unreasonable geometries (none were identified).

Further confidence in our models was built by comparison of the effects of S315T (the canonical mutation causing resistance) and S316G (a control mutation). The first is known to cause widespread resistance with little to no fitness cost [14,16,17,19], while the second which is located in the very next amino acid in the sequence is known not to cause resistance [26]. As we will see in the next sections, our models were able to predict this by demonstrating severe channel 1 constriction for S315T near the entrance to the catalytic heme, but show minimal constriction or even widening of the channels for S316G. (Tables 1 and 2) This provides further confidence in accuracy of our homology models and their ability to predict the phenotype.

### *3.2* Proximity of variants to Access Channels on M. tuberculosis-KatG

The amino acid substitutions were mapped on each of the 19 homology models generated,14 for novel mutations reported by Torres et al. [4] and five for control sequences. (These 19 structures are included as Supplementary Information.) Interestingly, the locations of the 14 amino acid substitutions appear to be distributed randomly throughout the protein sequence. (Figure 3) As the majority of these 14 amino acid substitutions are not found in or near the catalytic center, we hypothesized that perhaps the amino acid substitutions are causing steric constrictions in the access channels. For this reason, we decided to initially investigate the effect of the S315T substitution on the KatG tertiary structure.

### *3.3* Structural Origins of Resistance Due to the S315T Mutation

The S315T variant *M. tuberculosis*-KatG has been found in greater than 50% of clinical INH-resistant isolates and is the most prevalent INH resistance conferring mutation.[14,16,17,19] Using the x-ray crystal structures of WT- and S315T-*M. tuberculosis-KatG* proteins, 2CCA.pdb and 2CCD.pdb respectively, we measured the cross-sectional areas of a position near the surface as well as a position near the entrance to the catalytic heme in each of the three channels.

In Channel 1 of the S315T-*M tuberculosis*-KatG protein, a 22.11% reduction (as compared to the WT structure) in cross-sectional area and an obvious blockage in the path was observed near the entrance to the catalytic heme. (Table 2 and Figure 6) In addition, the hydroxyl group of the threonine side chain is positioned in the channel such that INH can readily interact with it. These observations are consistent with the widely held hypothesis that INH resistance originates in a constriction of this access channel in addition to hydrogen bonding to the bulkier sidechain of threonine.[14,16,17,19,20] In Channels 2 and 3 there were actually increases in crosssectional areas of 8.99% and 9.51% respectively.

**Figure 6.**
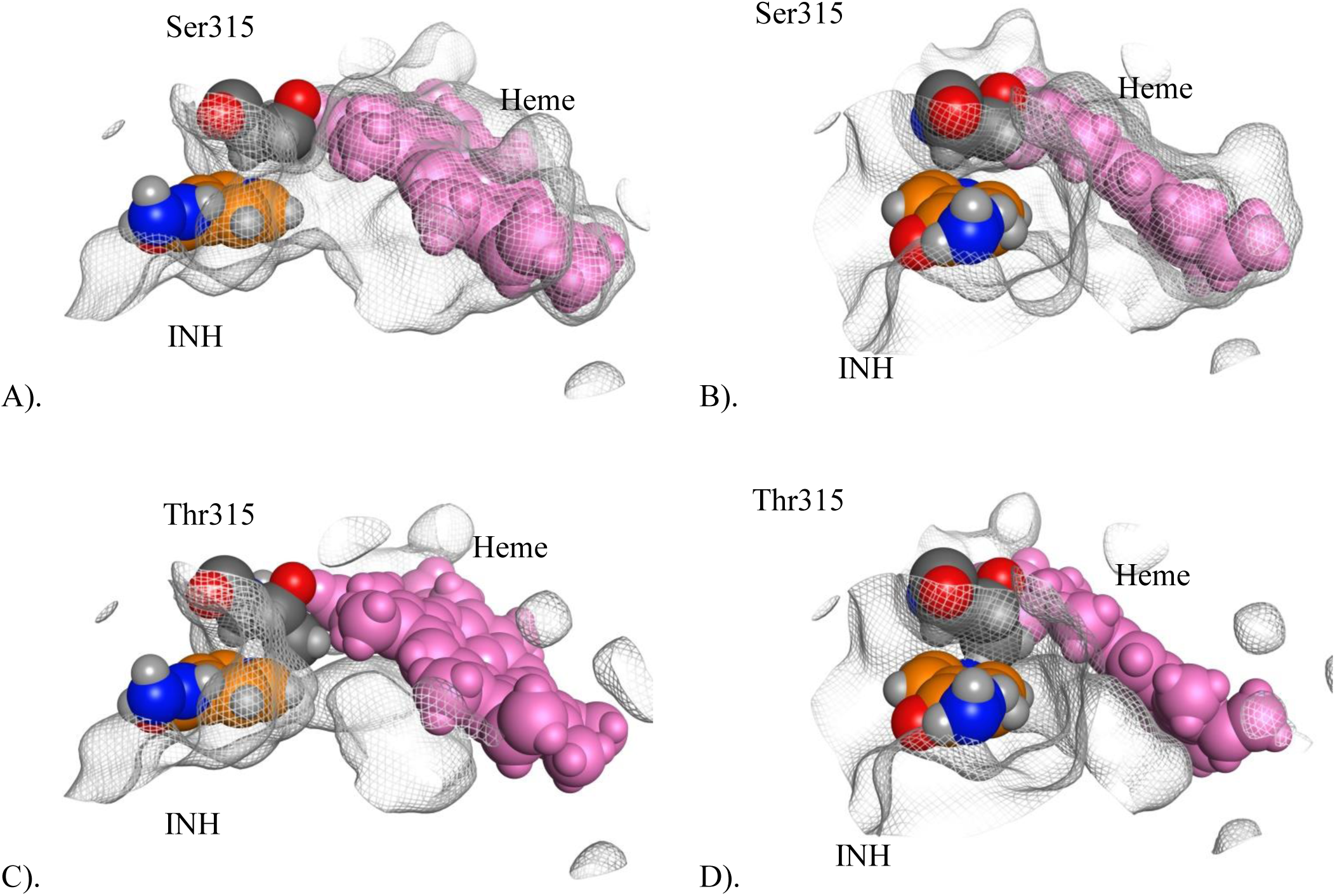
Illustration of the steric effect of S315T in Channel 1. Figures (A) and (B) show the configuration of the channel near the heme (pink) with Ser315 (dark grey) providing clear access to the heme for INH (orange). Figures (C) and (D) show the configuration of the channel near the heme with Thr315 (dark grey) and its larger side chain blocking the access channel, limited access to the heme. Horizontal (**A**) and head-on (**B**) screenshots obtained from the x-ray crystal structure of WT *M. tuberculosis*-KatG (2CCA.pdb) superimposed with INH bound in Channel 1 from WT *Se*-KatG (3WXO.pdb). A molecular surface was generated for 2CCA to illustrate there is an open path for INH to the catalytic heme. Horizontal (**C**) and head-on (**D**) screenshots obtained from the x-ray crystal structure of S315T *M. tuberculosis*-KatG (2CCD.pdb) superimposed with INH (in orange coloring) bound in Channel 1 from WT *Se*-KatG (3WXO.pdb). The heme is colored in pink and Thr315 in CPK coloring. A molecular surface was generated for 2CCD to illustrate there is *not* an open path for INH to the catalytic heme due to blockage from the side chain of Thr315.

In contrast, the surface entrances to each of the three channels of the S315T-M. *tuberculosis-KatG* protein all had relatively minor reductions in cross-sectional surface areas, between 2-5%. The roles of each of these three access channels are not well understood, but for this amino acid substitution, S315T, it appears that the constriction in Channel 1 near the entrance to the heme and the potential to hydrogen bond to the Thr sidechain is sufficient to convey resistance to INH while not seeming to affect KatG’s catalytic activity. The S315T amino acid substitution is therefore thought to be neutral with regards to fitness cost.[16] These results are consistent with the molecular modeling studies of Pimental et al, who studied additional mutations at this position (S315T, S315R, S315I, S315N, S315G).[22]

### *3.4* Structural Origins of Resistance Due to the 14 Novel M. tuberculosis-KatG Variants

As described above, homology models were generated for each of the 14 *M. tuberculosis-* KatG variant sequences and structurally superposed onto the *M. tuberculosis*-WT x-ray structure. The major and minor diagonals of each channel at points near the surface as well as near the catalytic heme were used to calculate the cross-sectional area of each channel. (ST1 and ST2) The calculated cross-sectional areas were compared to those of the WT KatG for each homology model. (Tables 1 and 2, Figures 4 and 5).

Table 1 lists the cross-sectional changes calculated near the surface entrance for each channel. The percent area change due to the novel mutations across the three channels ranged from −46.25% to −8.48% with mean cross-sectional area reductions of −19.47%, −27.96%, and − 30.79% for the three channels respectively.

Table 2 lists the cross-sectional changes calculated in each channel near the catalytic heme. For Channel 1, nine of the mutations only had minor reductions in cross-sectional areas ranging up to −7% and five actually had notable *increases* in cross-sectional areas resulting in a positive mean change in cross-sectional surface area of +2.03%. For Channel 2, all 14 variants had reductions in cross-sectional surface areas ranging between −9.85% and −21.06% with a mean *reduction* of −16.18%. For Channel 3 all 14 KatG variants had significant *increases* in cross-sectional areas ranging between +26.96 to +72.80% with a mean value of +52.50%.

In summary, while the 14 mutations had a variable effect on the opening near the heme, strikingly, all caused significant constriction of the opening of all three channels at the surface. These results are all consistent with our hypothesis that the 14 novel mutations cause synergistic constrictions in the channels such that INH’s access to the catalytic center and therefore its activation is significantly reduced (if not entirely blocked). Interestingly, some of these mutations which resulted in constrictions near the surface entrances of Channels 1 and 3 simultaneously resulted in *dilation* of the opening near the catalytic center. Channel 2, on the other hand, was constricted at both ends, by all 14 polymorphisms. Although it can be hypothesized that the constrictions near the surface are sufficient to restrict INH’s access, it is clear that there is a need for additional experimental studies.

We were also curious to see if we could find a correlation between the location of the amino acid substitution on the protein and its effect on the three channels. Although six (including S315T) are found in/close to a channel, nine of the 14 reported substitutions are located more than 10Å away from all channels (Tables 1 and 2). If our models and hypotheses are correct, then these act via long-range, perturbations of the KatG tertiary structure.

Finally, we were unable to identify a correlation between amino acid class/identity and the degree of cross-sectional area reduction in one or more of the channels.

### *3.5* Controls

In contrast to mutations whose causal role had been established by Torres et al., the control mutations caused much less constriction of the entrance to all three channels. The resulting change in the surface opening of Channel 1 ranged between −5.26% and +2.29%, with an average change of −1.11%. This is in sharp contrast with the effects of resistance-conferring mutations (minimum change: −8.48%, average change −19.47%). (Table 1)

The effects of the control mutations on Channels 2 and 3 surface openings were just as dramatically insignificant. Channel 2 changes due to the control mutations ranged between +0.09% and +4.80% with an average change of +1.98% as compared to the minimum change of −17.98% and average change of −27.96% for the resistance-causing group. In Channel 3, we observed a range of −2.26% to +4.43% changes with mean change of +1.11% for the control group in comparison to the minimum change of −10.96% and average change of −30.79% for the resistance-causing group.

Near the heme, the majority of control mutations had a widening effect. The resulting change in Channel 1 ranged between +9.98% and +14.50%, with an average change of +12.91%. Resistance causing mutations caused a variation between −6.95% and +25.77% with an average change +2.03%. (Table 2) For Channel 2 changes due to the control mutations ranged between −4.65% and +0.08% with an average change of −2.31% as compared to the minimum change of −9.85% and average change of −16.18% for the resistance-causing group. In Channel 3, we observed a range of +19.67% to +34.40% changes with mean change of +25.25% for the control group in comparison to the minimum change of +26.96% and average change of +52.50% for the resistance-causing group.

In summary, we observed a striking difference between the effects of the control mutations and the 14 novel resistance—causing mutations. In every case, control mutations either caused a widening or an insignificant amount of constriction on the channel openings. The 14 novel resistance causing mutations imposed a significant amount of constriction, especially at the opening near the surface. (Tables 1 and 2)

### *3.6* Fitness Costs

Significant constriction of the access channels can also impact the access of the substrate to the heme. As such, we hypothesize that those mutations that have significant constricting effects, especially on all three channels, would have a fitness cost. All 14 novel mutations fall in this category. The canonical S315T is an exception since it only constricts Channels 1 but it actually increases the diameter of Channels 2 and 3. Here, we put forth the hypothesis that perhaps it is because of this unusual steric effect that this mutation has such a low fitness cost and therefore is so prevalent among resistance conferring mutations.

### *3.7* R385W and D387G

These two mutations were first reported by Torres et al. [4] in clinical INH-resistant *M. tuberculosis* isolate that had no other mutations in KatG or promoter of *inhA*. Although isolates harboring the two mutations demonstrated very high resistance levels (R385W: MIC > 10) (D387G: 3 < MIC ≤ 10) in *M. tuberculosis*, mutagenesis in *M. smegmatis* failed to show a causal role in resistance. Our *M. tuberculosis* models demonstrate that both of these mutations cause significant constriction of the openings of the three channels near the surface in line with those associated with the models of few other novel mutations. (Tables 1 and 2). Perhaps this could be due to subtle differences between the steric effects in the two species. One hypothesis could be that these mutations restrict the access channels in a way that growth rate (as a result of substrate access restriction) is affected in *M. smegmatis* but not in *M. tuberculosis*. In such a case, *M. smegmatis* mutants with these variants would not reach the cut off growth units in a growth-based test and hence incorrectly be classified as susceptible while *M. tuberculosis* mutants would be able to do so and hence be identified as resistant. In order for this hypothesis to be true, such steric effects must be specific to these two mutations beyond what our models can demonstrate. Our modeling of *M. tuberculosis*-KatG variants is inconclusive and does not support this hypothesis since it shows that the constricting effects due to these mutations are significant on all three channels (Table 1) but are not the largest among the 14 mutations. So, it is difficult to explain why those with bigger constricting effects do not limit substrate access but do so in these two cases. As such, the role of these two mutations in resistance remains unclear. It is certainly possible that the two have no role in resistance and that an alternative mechanism elsewhere in the genome is in play.

### *3.8* Correlation of Steric Effects with Level of Resistance

Overall, with one notable exception of KatG-S315T model, the correlation of observed INH resistance with measurements of the surface channel cross-sectional areas were stronger than the correlations with measurements of the cross-sectional areas near the heme. Table 3 displays the Spearman correlation coefficients associated with 12 outstanding functions. Functions (1) through (3) are single variable functions and display the correlation between surface crosssectional areas of the three access channels (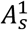,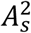,and 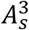) and MIC. Among these, the strongest correlation was observed with 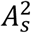, the surface cross sectional area of Channel 2, with *ρ_s_* = −0.82. The negative correlation indicates an inverse relationship (the smaller the surface opening of the channel, the higher the MIC value). Generally, the cross-sectional areas of the channels near the heme and the smallest cross-sectional area of each channel showed notably weaker correlations with MIC values. For Channel 2, for instance, functions (4) and (5) in Table 3, shows a weaker *ρ_s_* (-0.72) for the opening near the heme 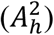, and the smallest cross-sectional area 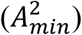.

**Table 3.** Spearman Correlation Coefficients between the proposed models’ outputs and MIC value. 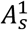, 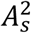, and 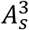 represent the cross-sectional areas of the openings of channels 1, 2, and 3, on the surface of the 3-D protein structure, respectively.

| Model | $\rho_s$ | Model | $\rho_s$ | Model | $\rho_s$ | Model | $\rho_s$ |
| --- | --- | --- | --- | --- | --- | --- | --- |
| (1) $A_s^1$ | -0.64 | (4) $A_h^2$ | -0.72 | (7) $A_{min}^2 + A_{min}^3 - A_{min}^1$ | +0.57 | (10) $A_s^2 - A_s^1$ | -0.51 |
| (2) $A_s^2$ | -0.82 | (5) $A_{min}^2$ | -0.72 | (8) $A_h^2 + A_h^3 - A_h^1$ | +0.38 | (11) $A_s^3 - A_s^1$ | +0.09 |
| (3) $A_s^3$ | -0.73 | (6) $A_s^2 + A_s^3 - A_s^1$ | -0.86 | (9) $A_s^2 + A_s^3 + A_s^1$ | -0.75 | (12) $A_s^2 + A_s^3$ | -0.81 |

Overall, among the 1,420 input-function combinations, the strongest correlation with MIC values was observed at the surface of the protein and is reported by function (6) in Table 3. The function is a linear combination of the surface openings of the three channels:

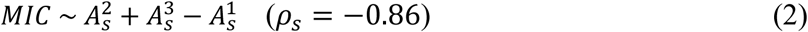

The counter parts of this function that took the minimum areas as input, function (7), and the areas near the heme, function (8), showed weaker correlations. (Table 3) All variations of this function, including the sum of the surface areas produced lower correlation—function (9) in the table. Dropping any channels from function (6), weakened the correlation—functions (10) through (12). In function (6), the negative *ρ_s_* value, positive coefficients of 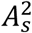 and 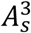, and a negative coefficient of 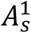 indicates that a reduction in the surface cross-sectional areas of channels 2 and 3 and an increase in the surface cross-sectional area of Channel 1 would increase the MIC value. While the roles of Channels 2 and 3 in increasing the MIC values was expected, the suggested role for Channel 1 was not and needs to be further investigated.

## 4 Conclusion

We have presented a structure-based hypothesis for the origin of INH resistance in 14 reported KatG variant proteins identified by Torres et al.[4] Our results show that resistance emerges due to constriction of one or more access channels from the KatG surface to the buried catalytic center. Our hypothesis is that these constrictions limit/prevent INH passage to the active site, thus preventing INH from becoming activated and inhibiting enzymes involved in synthesizing components of the *M. tuberculosis* cell wall.

Each homology model containing a reported novel amino acid substitutions had significant constriction in one or more channels while the controls did not. Statistical correlation indicates that the steric effects of the 14 novel mutations are different from that of the canonical S315T While the latter severely constricts Channel 1 near the heme, the former group severely constricts Channels 2 and 3 at the surface of the protein to cause increased MIC. Our results support the hypothesis that the 14 novel mutations constrict access channels broadly resulting in limited substrate access and a fitness cost. S315T does not constrict Channels 2 and 3 resulting in no fitness cost. This study provides the knowledge base for experimental work to confirm the molecular mechanism of INH resistance. We plan future studies for experimental validation of these in silico results.

---

**Supplementary Table ST1:** The dimensions and the narrowest cross-section (area) of the three access channels measured near the surface of the 3D structure of the protein. The homology model for each sequence is provided in a supplementary file named as listed in the last column of this Table. %Δ refers to percentage change of the cross-sectional area of the channel as compared to that of the WT channel. The WT and S315T structures, 2CCA.pdb and 2CCD.pdb respectively, were obtained from the RCSB Protein Data Bank (www.rcsb.org)

|  | Channel 1 |  |  |  | Channel 2 |  |  |  | Channel 3 |  |  |  |  |
| --- | --- | --- | --- | --- | --- | --- | --- | --- | --- | --- | --- | --- | --- |
|  | D1<br>(Å) | D2<br>(Å) | Area<br>(Å<sup>2</sup>) | %Δ | D1<br>(Å) | D2<br>(Å) | Area<br>(Å<sup>2</sup>) | %Δ | D1<br>(Å) | D2<br>(Å) | Area<br>(Å<sup>2</sup>) | %Δ | File Name |
| **WT** | 9.20 | 9.22 | 66.62 | 0.00 | 9.28 | 8.73 | 63.63 | 0.00 | 7.49 | 6.69 | 39.35 | 0.00 |  |
| **S315T** | 9.31 | 8.85 | 64.71 | -2.87 | 8.94 | 8.58 | 60.24 | -5.32 | 7.31 | 6.71 | 38.52 | -2.11 |  |
| **Novel mutations reported by Torres et al. [4]** |  |  |  |  |  |  |  |  |  |  |  |  |  |
| **Y64S** | 7.71 | 8.83 | 53.47 | -19.74 | 8.61 | 7.45 | 50.38 | -20.82 | 6.29 | 6.41 | 31.67 | -19.54 | Y64S.pdb |
| **Y95C** | 8.84 | 6.91 | 47.98 | -27.99 | 8.25 | 7.78 | 50.41 | -20.77 | 5.29 | 5.25 | 21.81 | -44.57 | Y95C.pdb |
| **P131T** | 8.8 | 7.95 | 54.95 | -17.52 | 8.33 | 7.36 | 48.15 | -24.32 | 6.36 | 6.44 | 32.17 | -18.26 | P131T.pdb |
| **A139P** | 10.00 | 7.17 | 56.31 | -15.47 | 7.33 | 7.24 | 41.68 | -34.49 | 5.16 | 5.22 | 21.15 | -46.25 | A139P.pdb |
| **D142G** | 8.91 | 7.02 | 49.13 | -26.26 | 7.51 | 8.79 | 51.85 | -18.52 | 5.42 | 5.31 | 22.60 | -42.56 | D142G.pdb |
| **A162V** | 7.94 | 8.24 | 51.39 | -22.87 | 7.05 | 7.30 | 40.42 | -36.47 | 6.26 | 6.44 | 31.66 | -19.55 | A162V.pdb |
| **G269D** | 8.69 | 8.15 | 55.62 | -16.51 | 7.06 | 7.21 | 39.98 | -37.17 | 6.22 | 6.28 | 30.68 | -22.05 | G269D.pdb |
| **T306P** | 8.79 | 7.80 | 53.85 | -19.17 | 8.63 | 7.70 | 52.19 | -17.98 | 5.37 | 5.51 | 23.24 | -40.95 | T306P.pdb |
| **R385W** | 9.04 | 8.25 | 58.57 | -12.08 | 8.17 | 7.28 | 46.71 | -26.58 | 6.61 | 6.75 | 35.04 | -10.96 | R385W.pdb |
| **D387G** | 8.70 | 7.56 | 51.66 | -22.46 | 8.38 | 7.41 | 48.77 | -23.35 | 5.78 | 5.89 | 26.74 | -32.06 | D387G.pdb |
| **T394M** | 8.63 | 8.21 | 55.65 | -16.47 | 7.60 | 7.16 | 42.74 | -32.83 | 5.68 | 5.85 | 26.10 | -33.69 | T394M.pdb |
| **Q439P** | 9.56 | 8.12 | 60.97 | -8.48 | 7.20 | 7.17 | 40.55 | -36.28 | 6.12 | 5.90 | 28.36 | -27.94 | Q439P.pdb |
| **F483L** | 8.85 | 7.27 | 50.53 | -24.15 | 8.12 | 7.40 | 47.19 | -25.83 | 5.23 | 5.98 | 24.56 | -37.58 | F483L.pdb |
| **A541D** | 9.25 | 7.02 | 51.00 | -23.45 | 7.19 | 7.21 | 40.71 | -36.01 | 5.77 | 5.63 | 25.51 | -35.17 | A541D.pdb |
| **Control Mutations** |  |  |  |  |  |  |  |  |  |  |  |  |  |
| **A110V** | 9.32 | 9.31 | 68.15 | +2.29 | 9.10 | 8.93 | 63.82 | +0.31 | 7.34 | 6.76 | 38.97 | -0.98 | A110V.pdb |
| **L499M** | 9.24 | 9.31 | 67.56 | +1.42 | 8.94 | 9.07 | 63.68 | +0.09 | 7.47 | 6.85 | 40.19 | +2.12 | L499M.pdb |
| **L587P** | 9.00 | 9.13 | 64.54 | -3.13 | 9.11 | 9.32 | 66.68 | +4.80 | 7.15 | 6.85 | 38.47 | -2.26 | L587P.pdb |
| **R463L** | 9.17 | 9.17 | 66.04 | -0.87 | 8.91 | 9.17 | 64.17 | +0.85 | 7.54 | 6.94 | 41.10 | +4.43 | R463L.pdb |
| **S316G** | 8.89 | 9.04 | 63.12 | -5.26 | 8.99 | 9.36 | 66.09 | +3.87 | 7.37 | 6.95 | 40.23 | +2.22 | S316G.pdb |

**Supplementary Table ST2:** The dimensions and the opening (area) of the three access channels measured near the heme at the catalytic center of the protein. The homology model for each sequence is provided in a supplementary file named as listed in the last column of this Table. %Δ refers to percentage change of the cross-sectional area of the channel as compared to that of the WT channel. The WT and S315T structures, 2CCA.pdb and 2CCD.pdb respectively, were obtained from the RCSB Protein Data Bank (www.rcsb.org).

|  | Channel 1 |  |  |  | Channel 2 |  |  |  | Channel 3 |  |  |  |  |
| --- | --- | --- | --- | --- | --- | --- | --- | --- | --- | --- | --- | --- | --- |
| | D1<br>(Å) | D2<br>(Å) | Area<br>(Å<sup>2</sup>) | %$\Delta$ | D1<br>(Å) | D2<br>(Å) | Area<br>(Å<sup>2</sup>) | %$\Delta$ | D1<br>(Å) | D2<br>(Å) | Area<br>(Å<sup>2</sup>) | %$\Delta$ | File Name |
| **WT** | 5.97 | 12.35 | 57.91 | 0.00 | 3.81 | 5.93 | 17.74 | 0.00 | 3.72 | 5.12 | 14.96 | 0.00 |  |
| **S315T** | 4.81 | 11.94 | 45.11 | -22.11 | 4.32 | 5.70 | 19.34 | +8.99 | 4.05 | 5.15 | 16.38 | +9.51 |  |
| **Novel mutations reported by Torres et al. [4]** |  |  |  |  |  |  |  |  |  |  |  |  |  |
| **Y64S** | 6.38 | 11.39 | 57.07 | -1.44 | 3.52 | 5.52 | 15.26 | -14.00 | 3.90 | 8.12 | 24.87 | +66.27 | Y64S.pdb |
| **Y95C** | 6.47 | 11.04 | 56.10 | -3.12 | 3.39 | 5.29 | 14.08 | -20.63 | 3.48 | 6.99 | 19.10 | +27.72 | Y95C.pdb |
| **P131T** | 6.34 | 11.46 | 57.06 | -1.46 | 3.57 | 5.40 | 15.14 | -14.67 | 3.89 | 8.10 | 24.75 | +65.43 | P131T.pdb |
| **A139P** | 8.12 | 11.42 | 72.83 | +25.77 | 3.34 | 5.34 | 14.01 | -21.06 | 3.43 | 7.05 | 18.99 | +26.96 | A139P.pdb |
| **D142G** | 6.74 | 11.19 | 59.24 | +2.29 | 3.43 | 5.39 | 14.52 | -18.17 | 3.73 | 7.50 | 21.97 | +46.88 | D142G.pdb |
| **A162V** | 5.94 | 11.55 | 53.88 | -6.95 | 3.38 | 5.99 | 15.90 | -10.39 | 3.79 | 7.78 | 23.16 | +54.81 | A162V.pdb |
| **G269D** | 6.64 | 11.52 | 60.08 | +3.75 | 3.40 | 5.72 | 15.27 | -13.92 | 3.70 | 7.69 | 22.35 | +49.39 | G269D.pdb |
| **T306P** | 6.36 | 11.50 | 57.44 | -0.80 | 3.47 | 5.48 | 14.93 | -15.84 | 3.86 | 7.85 | 23.80 | +59.09 | T306P.pdb |
| **R385W** | 6.46 | 11.62 | 58.96 | +1.81 | 3.41 | 5.46 | 14.62 | -17.59 | 3.74 | 7.79 | 22.88 | +52.97 | R385W.pdb |
| **D387G** | 6.46 | 11.12 | 56.42 | -2.57 | 3.40 | 5.32 | 14.21 | -19.94 | 3.69 | 7.83 | 22.69 | +51.70 | D387G.pdb |
| **T394M** | 6.33 | 11.54 | 57.37 | -0.92 | 3.52 | 5.75 | 15.90 | -10.42 | 3.94 | 7.89 | 24.42 | +63.22 | T394M.pdb |
| **Q439P** | 7.31 | 11.69 | 67.12 | +15.90 | 3.67 | 5.55 | 16.00 | -9.85 | 3.97 | 8.29 | 25.85 | +72.80 | Q439P.pdb |
| **F483L** | 6.33 | 11.20 | 55.68 | -3.84 | 3.37 | 5.31 | 14.05 | -20.80 | 3.56 | 7.53 | 21.05 | +40.74 | F483L.pdb |
| **A541D** | 6.60 | 11.17 | 57.90 | -0.01 | 3.41 | 5.35 | 14.33 | -19.25 | 3.89 | 7.69 | 23.49 | +57.06 | A541D.pdb |
| **Control Mutations** |  |  |  |  |  |  |  |  |  |  |  |  |  |
| **A110V** | 6.77 | 12.47 | 66.30 | +14.50 | 3.82 | 5.73 | 17.19 | -3.12 | 3.74 | 6.12 | 17.98 | +20.17 | A110V.pdb |
| **L499M** | 6.77 | 12.49 | 66.41 | +14.69 | 3.88 | 5.72 | 17.43 | -1.77 | 3.70 | 6.16 | 17.90 | +19.67 | L499M.pdb |
| **L587P** | 6.55 | 12.38 | 63.69 | +9.98 | 3.96 | 5.71 | 17.76 | +0.08 | 3.62 | 6.35 | 18.05 | 20.69 | L587P.pdb |
| **R463L** | 6.75 | 12.37 | 65.58 | +13.25 | 3.86 | 5.73 | 17.37 | -2.10 | 3.75 | 6.67 | 19.64 | +31.32 | R463L.pdb |
| **S316G** | 6.72 | 12.30 | 64.92 | +12.11 | 3.84 | 5.61 | 16.92 | -4.65 | 3.77 | 6.79 | 20.10 | +34.40 | S316G.pdb |

**Supplemental Table ST3:** Functions used in estimation of MIC values

| Statistical | Polynomial | Rational | Exponential |
| --- | --- | --- | --- |
| $f(x) = stdev(x)$ | $f(x) = x$ | $f(x, y, z) = \frac{x}{y + z}$ | $f(x) = x^2$ |
| $f(x, y, z) = max(x, y, z)$ | $f(x, y) = x + y$ | $f(x, y, z) = \frac{x + y}{z}$ | $f(x) = \sqrt{x}$ |
| $f(x, y, z) = min(x, y, z)$ | $f(x, y) = x - y$ | $f(x, y) = \frac{x}{y}$ | $f(x) = x^3$ |
| | $f(x, y, z) = x + y + z$ | $f(x, y, z) = \frac{x}{y \times z}$ | $f(x) = \sqrt[3]{x}$ |
| | $f(x, y, z) = x + y - z$ | $f(x, y, z) = \frac{x^2}{y \times z}$ | |
| | $f(x, y, z) = x - y - z$ | | |

**Note:** $x$ , $y$ and $z$ could be any of $A_s$ , $\% \Delta A_s$ , $A_h$ , $\% \Delta A_h$ , and $A_{min}$ values for any of the Channels 1, 2, and 3. For instance $f(x, y, z) = \frac{x+y}{z}$ , represent all variations of the five measurements from the three channels such as $f(A_s^1, A_s^2, A_s^3) = \frac{A_s^1 + A_s^2}{A_s^3}$ , $f(A_s^1, A_s^2, A_s^3) = \frac{A_s^3 + A_s^2}{A_s^1}$ , $f(\% \Delta A_s^1, \% \Delta A_s^2, \% \Delta A_s^3) = \frac{\% \Delta A_s^1 + \% \Delta A_s^3}{\% \Delta A_s^2}$ , etc., where $A_s^1$ , for instance, is the cross-sectional area of Channel 1 measurement at the surface of the protein structure.

**Supplementary Figure SF1.**
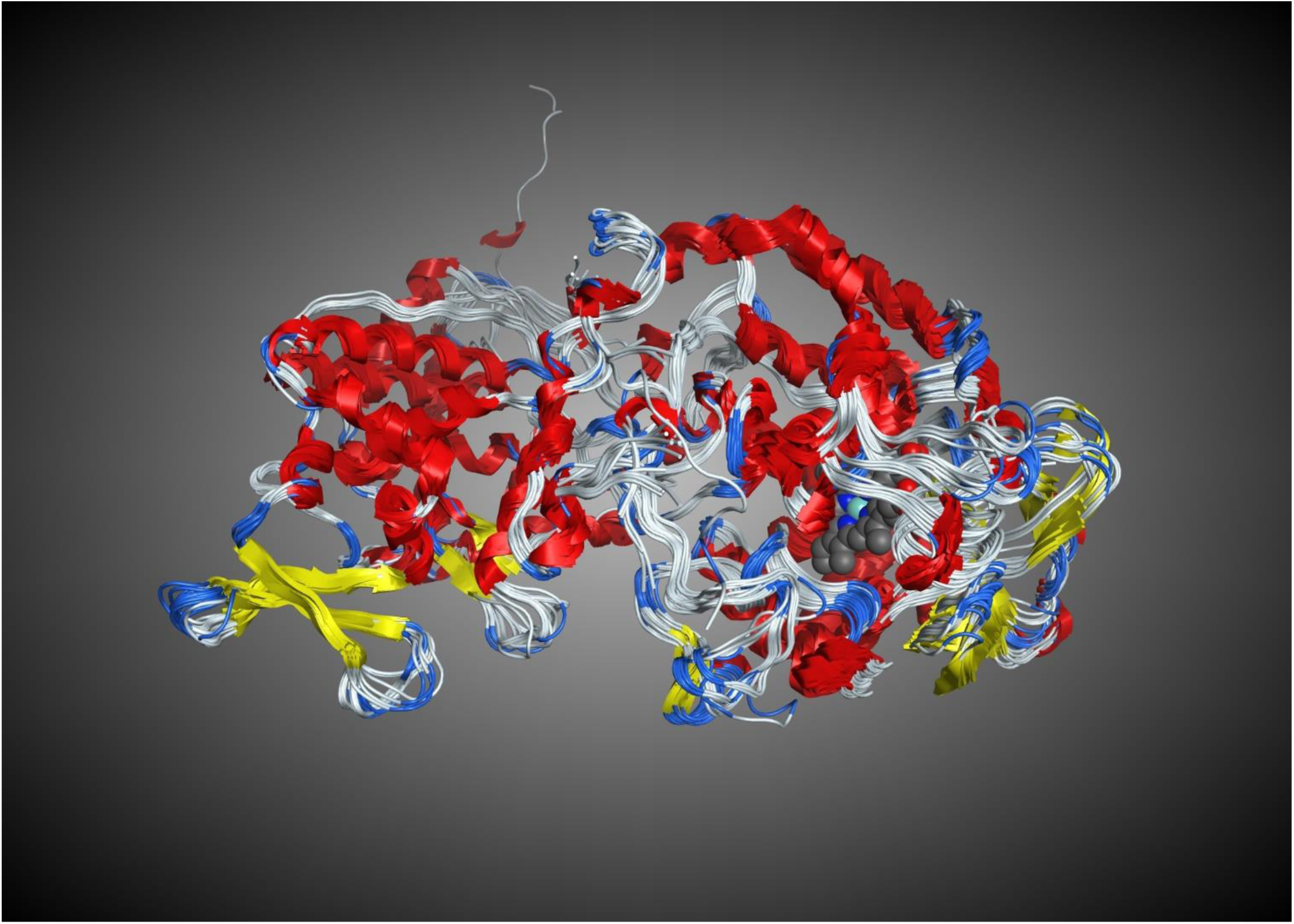
Structural superposition of the 48 x-ray crystal structures of KatG from six species illustrating the similar secondary and tertiary structures, overall 0.91 Å *RMSD*. The heme from 2CCA.pdb is shown in blue and grey.

